# Opposing regulation by Rev1 of DNA polymerase zeta activity on damaged versus undamaged DNA

**DOI:** 10.64898/2026.01.21.700666

**Authors:** Rachel Bezalel-Buch, Carrie Stith, Alena V. Makarova, Sara K. Binz, Peter M. Burgers

**Author notes:** AVM. Institute of Gene Biology, Russian Academy of Sciences, Moscow, 119334, Russia; SKB, Confluence Discovery Technologies, St. Louis, MO, USA.

## Abstract

The Rev1 deoxycytidyl transferase functions as a scaffold protein for DNA polymerase ζ (Pol ζ)-mediated translesion synthesis (TLS). Biochemical studies with yeast enzymes indicate that Rev1 plays a dual regulatory role in TLS, stimulating Pol ζ activity at sites of damage but inhibiting its activity on undamaged DNA. An evolutionary conserved N-terminal alpha-helical motif (M1), located 10-20 amino acids upstream of Rev1’s single BRCT domain, is required for the inhibitory activity of Rev1 on undamaged DNA. Mutations in the M1 motif result in a stimulation of Pol ζ replication activity on both undamaged and damaged DNA. Yeast cells carrying a *REV1* mutant lacking the M1 motif, show a four-fold increase in complex mutations, without significantly affecting overall spontaneous mutation rates. A catalytically inactive mutant of Rev1 still exerts these regulatory functions. However, regulation requires that Rev1 and Pol ζ form a stable complex, and that this complex is coordinated by the replication clamp PCNA.

**Graphical Abstract:** 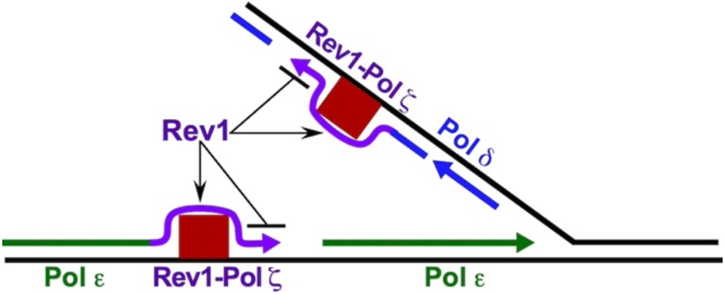

Rev1 stimulates lesion bypass by Pol ς but inhibits Pol ς activity on undamaged DNA

## INTRODUCTION

The eukaryotic genome is constantly exposed to different exogenous and endogenous DNA-damaging factors that threaten its integrity (1,2). The damaged DNA (e.g, nicks and breaks, base loss, base alteration, and interstrand crosslinks) is being repaired by a plethora of DNA repair mechanisms that exist in the cell, such as Homologous Recombination, Non-Homologous End-Joining, Nucleotide Excision Repair, Base Excision Repair, and Mismatch Repair (3-7). Despite the variety of repair mechanisms, the replicative polymerases will likely encounter a lesion within the DNA template during each cell cycle. This could result in stalling of DNA synthesis (8-10). To complete the synthesis of the genome before mitosis and to avoid the deleterious consequences of prolonged replication fork stalling, cells adopt DNA Damage Tolerance (DDT) pathways that allow the replication machinery to proceed without repairing the damaged DNA (11,12). This pathway has two branches: error-free and error-prone (13,14). The error-prone DDT pathway consists of translesion synthesis (TLS) (15,16), which is operated by low-fidelity and low processivity TLS DNA polymerases, using a two-step mechanism – mis-incorporation across damage, followed by extension of the resulting, distorted template/primer (14,15,17,18). TLS mechanisms and machineries are conserved through evolution and include DNA polymerase ζ (Pol ζ) and members of the Y-family polymerases, which include Pol η, Pol κ, Pol ι, and Rev1 in human, and Pol η and Rev1 in *S. cerevisiae* (19).

Pol ζ is a multi-subunit complex, composed of the catalytic subunit Rev3 and four additional accessory subunits (2 copies of Rev7, which form a dimer, Pol31, and Pol32) assembled into a ring-like structure (Figure 1A) (20-22). Rev3 belongs to the B-family of DNA polymerases (23-25). It shares Pol 31 and Pol 32 with the replicative DNA polymerase δ (26-28), but lacks 3’ to 5’ exonuclease activity. Rev3 interacts with Rev7 and Pol31, and Pol32 interacts with Pol31 and Rev7 (Figure 1A) (22). All of the accessory subunits are crucial for complex stability (26). Pol ζ is considered to play a role in the extension step of lesion bypass (29).

**Figure 1.**
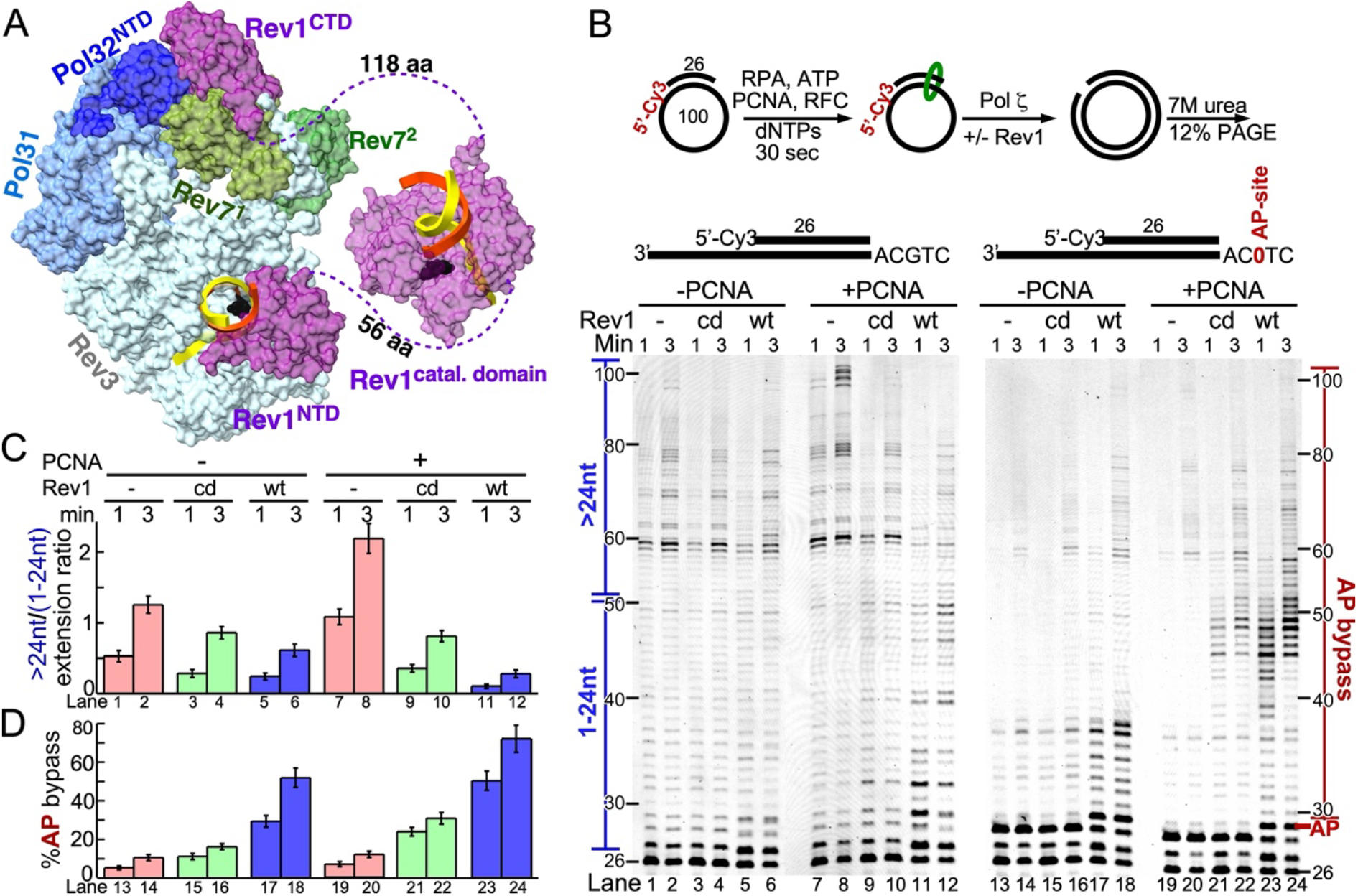
Rev1 plays a key role in Pol ζ regulation. (A) CryoEM structure of Rev1-Pol ζ (PDB: 8tlq) at 3.5 Å resolution. The catalytic domain of Rev1 is unresolved in the cryoEM structure but has been added from the crystal structure (PDB: 6×70; (66)) for illustrative purposes. Note that Pol ζ has two copies of the Rev7 subunit, shown in different shades of green. (B) Top, scheme of the assay. 7 M urea 12% PAGE analysis of replication reactions by Pol ζ in the absence or presence of PCNA, and either without Rev1 or wild-type (wt) or catalytic dead (cd) Rev1. The assays in lanes 1-12 were performed with undamaged DNA, whereas the assays in lanes 13-24 were performed on a template-primer carrying a model abasic (tetrahydrofuran) site, as shown above the gel. On the left side of the gel is indicated what we define as short versus long products used in the calculation in (C). On the right side of the gel is indicated what is considered AP-site bypass (D). (C-D) Quantification of the data in (B). Plotted is the ratio of long over short extension products as a measure of extension efficiency for lanes 1-12 (top graph), and the percentage of AP bypass for lanes 13-24 (bottom graph).

Rev1 consists of an N-terminal BRCT domain for binding PCNA and DNA (30-32), a motif for binding other Y-family DNA polymerases, ubiquitin-binding motifs (UBM), and a C-terminal interaction domain with Rev7 and Rev3 (33,34). In addition, Rev1 has two PCNA-binding regions, one localized to the BRCT domain (30,35), and a second one in the PAD (polymerase-associated domain), also called little finger domain of the catalytic module (36-38). Rev1 has an unique deoxycytidyl (dCMP) transferase activity (39), inserting dCMPs rather indiscriminately, but with highest activity across abasic sites (AP) and template G residues (40-42). Rev1’s catalytic activity is biologically relevant for some DNA lesions such as AP-sites and 1,N^6^-ethenoadenine (43,44), but not necessary for the majority of TLS and mutagenesis events. The efficiency of mutagenesis with the catalytic-dead Rev1 and the mutation spectrum is affected to varying degrees, depending on the type of DNA damage (45). In addition, beyond Pol ζ, other DNA polymerases, such as Pol η or Pol δ in yeast, may participate in the bypass of damage in Rev1 catalytic activity-compromised cells (46-48).

Thus, the primary function of Rev1 is considered that of a scaffold protein for assembly and function of the mutasome complex (34,46,49). With few exceptions (50,51), most previous biochemical studies of TLS have focused either on Rev1 or on Pol ζ, but not on the integrated complex together with the PCNA clamp. In this study, we have explored the regulatory function of Rev1 with regards to Pol ζ activity, depending on the state of the DNA, whether damaged or undamaged, dNTP availability, and the presence of PCNA. We show that Rev1 protein has a dual function in that it stimulates bypass by Pol ζ at sites of damage but inhibits Pol ζ activity on undamaged DNA. We identified a motif (M1) upstream of the BRCT module (22), which is required for the inhibitory function of Rev1 on undamaged DNA, and describe the consequences of mutation in the M1 motif for mutagenesis.

## MATERIAL AND METHODS

### Reagents and enzymes

Chemical reagents were purchased from Sigma-Aldrich (St. Louis, MO, USA). USER™ Mix, T4 polynucleotide kinase, and T4 DNA ligase were purchased from New England Biolabs (Ipswich, MA, USA). Pfu Turbo DNA polymerase was purchased from Agilent. dATP, dCTP, dTTP, and dGTP were purchased from Invitrogen. ZR small-RNA PAGE recovery kit was purchased from Zymo Research. Plasmid purification kit and PCR purification/Gel extraction kit were purchased from Qiagen.

### Oligonucleotides and primers

All oligos were purchased from IDT and were purified by either polyacrylamide gel electrophoresis or high-pressure liquid chromatography. The model abasic site used in all TLS experiments (AP1) is a tetrahydrofuran moiety, designated as /idSP/ by IDT. The sequences of the templates and primers are in Supplementary Table S2.

### Preparation of 100-mer circular ssDNA substrates

The 100-mer sequence was derived from Bluescript SK2 DNA. To create 100-mer circular ssDNA, the SKII-100 and SKII-99AP1 oligos were hybridized to the bridging primer SKII 100 Anneal (Supplementary Table S2). T4 DNA Ligase was added to the reaction and incubated at 16°C overnight. Reactions were quenched with 50% formamide, 10 mM EDTA, and 0.1% SDS and loaded onto a 12% polyacrylamide-7 M urea preparative gel. The gel was stained with SYBR Safe and visualized on a SmartBlue Transilluminator (ACCURIS Instruments). The circular DNA substrate was isolated from the gel by using the ZR small RNA PAGE Recovery Kit (Catalog #R1070 – Zymo Research).

### *S. cerevisiae* strains and plasmids

The haploid strain PY435 (MATa *his3Δ-1 leu2-3,112 trp1 ura3-52 pep4Δ::HIS3 nam7Δ::KanMX4 rev1Δ::HYG rev3Δ::NAT*) was made by transforming strain PY330 (51) with a *rev3Δ::NAT* cassette, and was used to overproduce Pol ζ and Rev1. The two plasmids used for overproduction of Pol ζ were pBL818 (2 μM ori, *URA3, GAL1*-ZZtag-*REV3, GAL10*-*REV7*) and pBL347-TRP (2 μM ori, *TRP1, GAL1-POL31, GAL10-HIS*_*7*_*-POL32*), as described before (26), with a slight modification in which pBL347-TRP (with a *TRP1* selection cassette) replaced pBL347 (with a *LEU2* selection cassette). For overexpression, *REV1* and *REV1* mutants were cloned into pRS424-ZZtag (2 μM ori, *TRP1, GAL10-ZZtag*) using the Sal1 restriction site and named pBL827 (2 μM ori,*TRP1, GAL10*-ZZtag*-REV1*). The ZZ-tag, used in affinity purification on IgG beads, is separated from *REV1* by a HRV-3C protease site. pBL827 was used to overexpress Rev1 wild-type and mutants in yeast. pBL829 (pBluescript SK2, *CEN ARS HIS3 REV1*) was used for complementation analysis with *REV1* expressed under its native promoter. Specific primers were designed to generate the mutants listed in Supplementary Table S1. pBL827 and pBL829 were used as templates for site-directed mutagenesis. The site-directed mutagenesis protocol, using Pfu-turbo DNA polymerase, was according to the manufacturer (Agilent). All plasmid sequences were confirmed by sequencing. The Rev1 mutants used are listed in Supplementary Table S1. For spontaneous mutation and UV mutagenesis studies, we used haploid strains that were derived from the wild-type strain CL1265-7C (47), which was made Arg^+^ (from *arg4-17*) by transformation with the wild-type *ARG4* gene: PY452 (MATa *his3Δ-1 leu2-3,112 trp1 ura3-52*). Derived from PY452 by transformation with deletion cassettes, are PY476 (as PY452, *rev1Δ::NAT*), PY480 (as PY452, *rev3Δ::HYG*), and PY481(as PY452, *rev1Δ::NAT rev3Δ::HYG*).

### Protein expression and purification

*S. cerevisiae* replication protein A (RPA), replication factor C with a N-terminal truncation (Rfc1-Δ273) (RFC), and PCNA of were expressed and purified from *E. coli* (52-56). Pol ζ (Rev3–Rev7×2–Pol31–Pol32) and Rev1^wt^ and Rev1 mutants were overexpressed and purified as described before (26,50,51), with some changes. To avoid contamination of overproduced Rev1 with cellular Pol ζ and overproduced Pol ζ with cellular, wild-type Rev1, we used strain PY435 deleted for both *REV1* and *REV3*. Galactose induction was carried out for 16 hours to overexpress Pol ζ and for 4 hours to overexpress Rev1. Pol ζ was purified as described previously in (26).

Purifications of Rev1 wild-type and Rev1 mutants are based on ∼50 g of cells. Ammonium sulfate precipitation pellets were re-suspended in buffer R_0_ (50 mM Hepes (pH 8.0), 8% glycerol, 0.1% Tween 20, 0.01% C_12_E_10_, 1 mM DTT, 10 µM pepstatin A, 10 µM leupeptin, 2.5 mM benzamidine, 0.5 mM PMSF and 1 mM EDTA) and gently agitated with 1.0 ml of IgG Sepharose beads (GE Healthcare) for 2 hours. The beads were batch-washed with 20-bed volumes of Buffer R_eq_ (Buffer R_0_ + 500 mM NaCl), then batch-washed with 30-bed volumes of R_0.7_ (Buffer R_0_ + 700 mM NaCl), followed by resuspension in 20-bed volumes of R_0.7_ and packed into a disposable 20-ml BioRad column. Beads were washed extensively with 100-bed volumes of Buffer R_ATP_ (Buffer R_0_ + 400 mM NaCl, 1 mM ATP, 8 mM Mg acetate), followed by a wash with 20-bed volumes of Buffer R_0.2_ (Buffer R_0_ + 200 mM NaCl). The beads were resuspended in 10-bed volumes of Buffer R_0.2_ with PreScission protease and digested overnight at 4 °C with gentle rotation of the capped column. After collecting the eluent, the beads were washed with an additional 4-bed volumes of Buffer R_0.2_. Fractions were combined, diluted with buffer to a conductivity equivalent to 100mM NaCl, and loaded onto a 1 ml Heparin column that was pre-equilibrated and washed with 10-bed volumes of Buffer B1 (50 mM Hepes (pH 8.0), 100 mM NaCl, 8% glycerol, 0.01% C_12_E_10_, 1 mM DTT), followed by a wash with 20-bed volumes of Buffer B2 (50 mM Hepes (pH 8.0), 200 mM NaCl, 8% glycerol, 0.01% C_12_E_10_, 1 mM DTT). The column was eluted with 3-bed volume of Elution Buffer (50 mM Hepes (pH 8.0), 0.7M NaCl, 8% glycerol, 0.01% C_12_E_10_, 1 mM DTT, 0.1% Ampholytes pH3-10). Fractions were dialyzed against Buffer D (50 mM Hepes (pH 8.0), 200 mM NaCl, 8% glycerol, 0.01% C_12_E_10_, and 1 mM DTT, 0.1% Ampholytes) and stored at -80 °C.

### DNA polymerase activity assay

To reconstitute the Rev1-Pol ζ complex, Rev1 and Pol ζ were premixed in a 1.5:1 molar ratio on ice. DNA replication and primer extension assays were performed as described previously (51). The10 μl assays contained 40 mM Tris–HCl pH 7.8, 1 mM DTT, 0.2 mg/ml bovine serum albumin, 8 mM Mg acetate, 125 mM NaCl, 0.5 mM ATP, damage-response concentrations of dNTPs: 195 μM dCTP, 383 μM dTTP, 194 μM dATP, 50 μM dGTP (51,57), and DNA and factors/enzymes. A 5’-Cy3 26-mer primer (Table S2) was hybridized to the 100-mer circular undamaged (SKII-100) or damaged DNA (SKII 99-mer AP1). 10 nM primer/template in assay mix was coated with 40 nM RPA, and 30 nM PCNA was loaded by 10 nM RFC for 30 sec at 30°C. The reactions were initiated with 10-25 nM of Pol ζ or Rev1-Pol ζ. Reactions were incubated at 30°C for 0.5-3 minutes. Reactions were stopped with 50% formamide, 10 mM EDTA, and 0.1% SDS and analyzed on a 12% polyacrylamide-7 M urea gel. Visualization was done by fluorescence imaging on a Typhoon system and quantified by ImageQuant. All experiments were carried out at least in duplicate, and most were carried out 3-5 times. The values in Figures 1 and 2 are the mean and standard deviation from two independent replicates.

**Figure 2.**
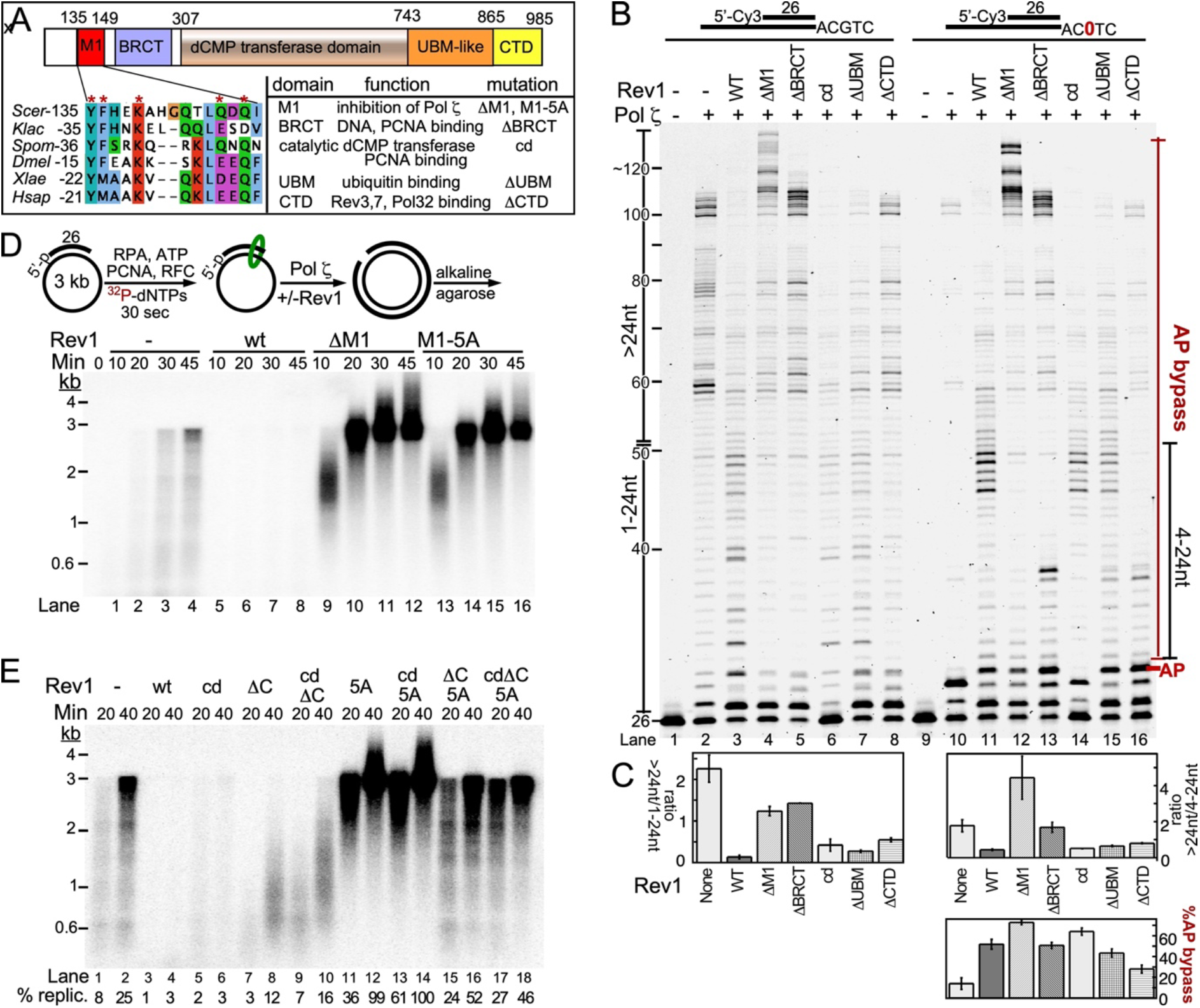
Rev1 domain analysis. (A) Rev1 protein domain map with below it a Multiple Sequence Alignment of the 15 amino acids M1 sequence from *Saccharomyces cerevisiae, Kluyveromyces lactis, Schizosaccharomyces pombe, Drosophila melanogastor, Xenopus laevis* and *Homo sapiens*. (B) 12% UREA PAGE of primer extension, DNA substrate used in the assay were SKII-100/SKII-AP1 circular ssDNA primed with SKII primer 1,(lanes 1-8) and bypass assay (lanes9-16). On the left side of the gel is indicated what we define short versus long products used in the calculation below (C). On the right side of the gel is indicated what is considered AP-bypass, and what are short extension products (4-24 nt) after AP-bypass. (C) Quantification of the data in (B), similar to Figure 1B,C. (D) Replication of primed 3 kb Bluescript SKII ssDNA. Top, scheme of the assay. Assays were carried out for increasing times with the indicated factors, and the products were separated on a 1% alkaline agarose gel.(E). Similar assay with mutant forms of Rev1 that combine domain mutants.

### Nucleotide omission assay

The primer extension assay has been slightly modified. In a 10 μl reaction, the mix included 40 mM Tris–HCl pH 7.8, 1 mM DTT, 0.2 mg/ml bovine serum albumin, 8 mM Mg acetate, 125 mM NaCl, 0.1 mM AMP-CPP, 10 nM 5’-Cy3-primed 100-mer circular ssDNA template. The DNA was coated with 40 nM RPA and 30 nM PCNA was loaded by 10 nM RFC and polymerases were bound with 25 nM of Pol ζ or Rev1-Pol ζ (1.5:1 molar ratio) for 30 sec at 30 °C. AMP-CPP (α,β-methyleneadenosine 5′-triphosphate) (0.5 mM) was included in this assay as an active ATP analog for PCNA loading, but inactive as a nucleotide precursor for the DNA polymerase. This avoids complications with the use of ATP as a PCNA loader, since ATP contains minute contaminations of dATP (58). DNA synthesis was initiated with 10 μM of each of the four dNTPs or a dNTP mix lacking one nucleotide. This low dNTP concentration was used to minimize cross-contamination. Controls included Rev1 only reactions in Figure S6. Reactions were incubated at 30°C for 10 min and were stopped with 50% formamide, 10 mM EDTA, and 0.1% SDS and analyzed on a 12% polyacrylamide-7 M urea gel. Visualization was done by fluorescence imaging on a Typhoon system. The dNTP omission assays were carried out four independent times, with slight variations (being with or without specific Rev1 mutants), with one experiment reported in Figure 4 and a second experiment in Figure S6.

### Measurement of DNA polymerase ζ processivity in vitro

The standard 10 μl DNA polymerase assays contained 10 or 15 nM of primed (^32^P-SKII primer 4) circular SK2-100 ssDNA, coated with 40 nM RPA. PCNA (30 nM) was loaded with 10 nM RFC for 30 seconds at 30°C. The reaction was started by adding 2 or 3 nM, respectively of Pol ζ, Rev1-Pol ζ, or Rev1^ΔM1^-Pol ζ. After 0.5, 1, 1.5, and 2 min at 30°C, reactions were quenched with 50% formamide, 10 mM EDTA, and 0.1% SDS. The products were analyzed on 12% polyacrylamide-7 M urea gel. Quantification was done by phosphorimaging of the dried gel (^32^P). Plot profiles of all lanes were generated using ImageJ (after linearization of the gel data). These profiles plotted the intensity signal in the gel against an arbitrary y-coordinate. After background subtraction, we determined the y-coordinate and thereby the nucleotide position at which the median of the total signal of extended products was located (58). The processivity assays were carried out three independent times, with one experiment reported in Fig. 3 and a second experiment in Fig. S5.

**Figure 3.**
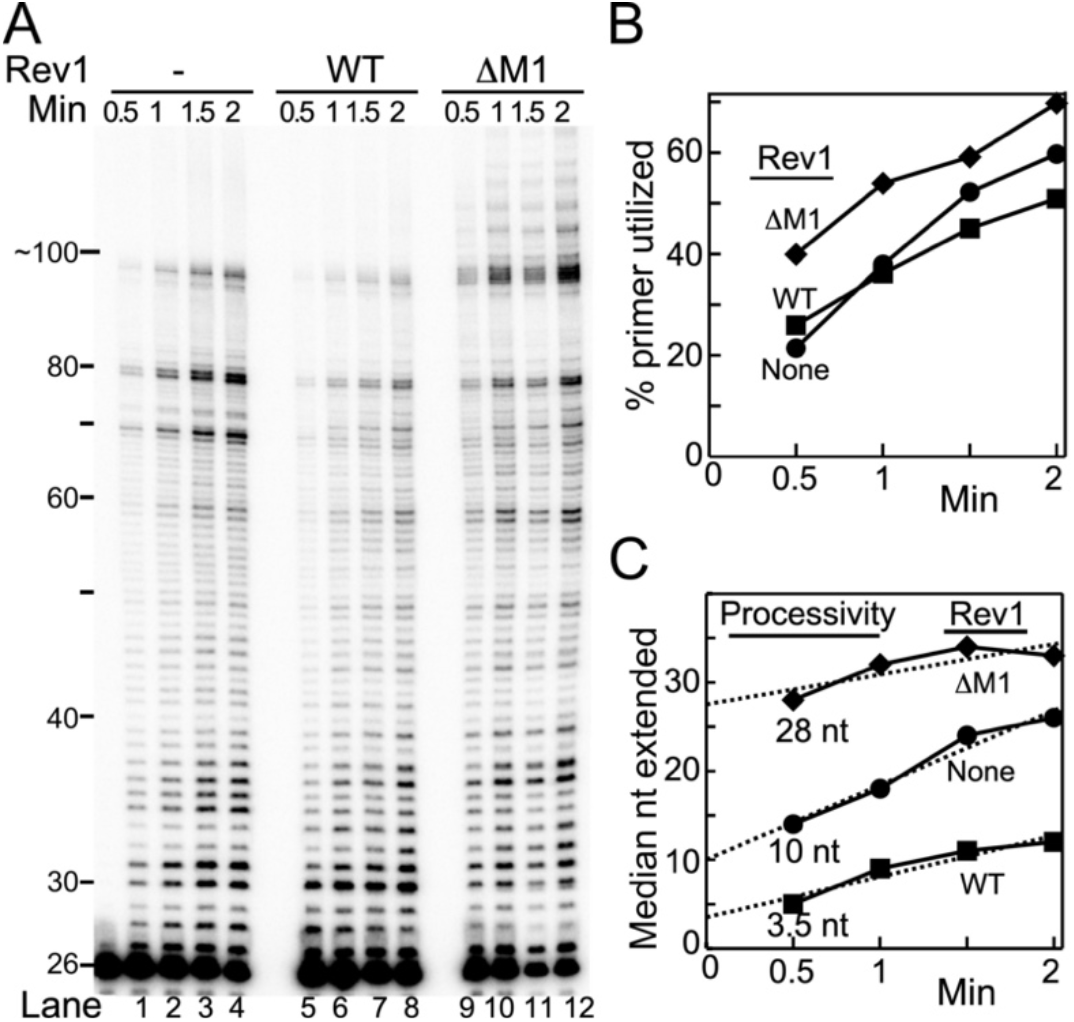
Rev1 determines Pol ζ replication processivity. (A) 12% UREA-PAGE analysis of a time-course replication assay containing 3 nM Pol ζ, or Rev1^wt^-Pol ζ or Rev1^ΔM1^-Pol ζ, and 15 nM of the DNA substrate (DNA substrate used in the assay was SKII-100 circular ssDNA primed with 5’-^32^P-labeled SKII primer 4). (B) Plot of the % primer utilized with time. (C) The median nucleotide positions, of extended products, was calculated from the intensity traces of each time point and plotted against reaction time. Linear plots were extrapolated to time zero to determine processivity.

**Figure 4.**
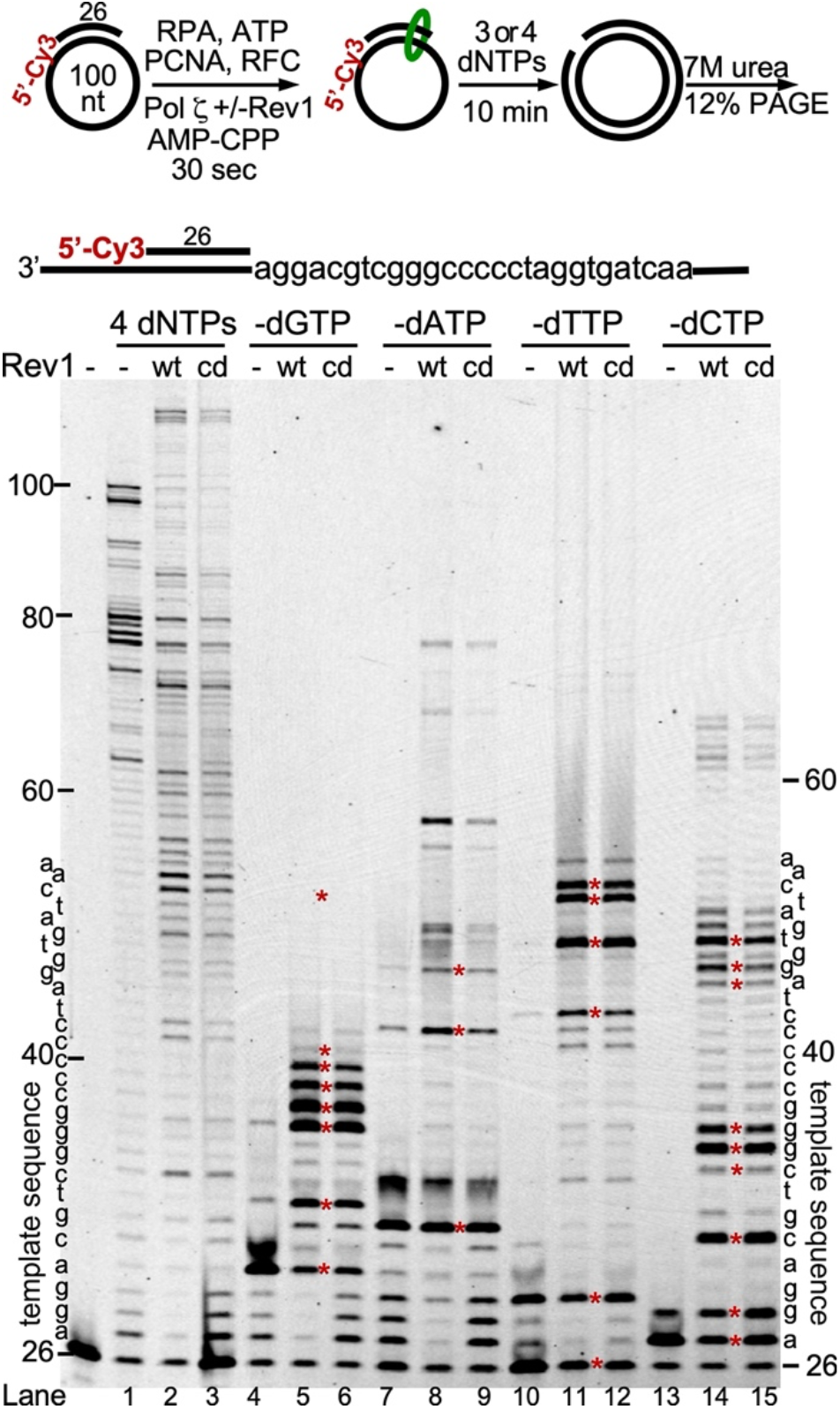
Rev1 stimulates Pol ζ infidelity upon dNTP omission. Top, diagram of assay and template sequence (DNA substrate used in the assay was SKII-100 circular ssDNA primed with SKII primer 3). dNTPs were present at 10 μM each and the reaction was terminated after 10 min and analyzed by 7 M Urea-12% PAGE. The template sequence is indicated on the sides of the gel. The red stars represent stall positions; each of these positions is located one nucleotide before the template nucleotide that is paired with the missing dNTP in the assay. For AMP-CPP usage, see Materials and Methods. Note that the replication data with 4 dNTPs (lanes 1-3) cannot be compared to those in Figures 1 and 2, because the assay conditions are very different (different primer, different dNTP concentrations, assay times).

### Spontaneous and damage-induced mutagenesis

The strains analyzed were PY452 (wild-type), PY476 (*rev1Δ*), PY480 (*rev3Δ*), and PY481(as PY452, *rev1Δ rev3Δ*). PY476 (*rev1Δ*) was transformed with pRS313 (Bluescript SK2, *CEN ARS HIS3)*, pBL829-0 (pBL313, *REV1*), or pBL829-M1 (pBL313, *rev1-ΔM1*). Fluctuation analyses to determine spontaneous mutation rates were carried out in duplicate. For each strain, 15 or 17 independent cultures were started at ∼1000 cells/ml and grown to saturation at 30 °C in minimal complete media (SC), and in minimal complete, selective media (SC-His) for analyses of *rev1-Δ* strains containing pBL829 plasmids. Appropriate dilutions were plated on SC-His medium, lacking arginine and containing 80 mg/liter of canavanine for mutant counting, and on SC-His-Arg medium for viable cell counting. Mutation rates were calculated from mutation frequencies using the method of the median according to Lea and Coulson (59), with the 95% confidence limits determined by standard statistical methodology (Figure 5A).

**Fig 5.**
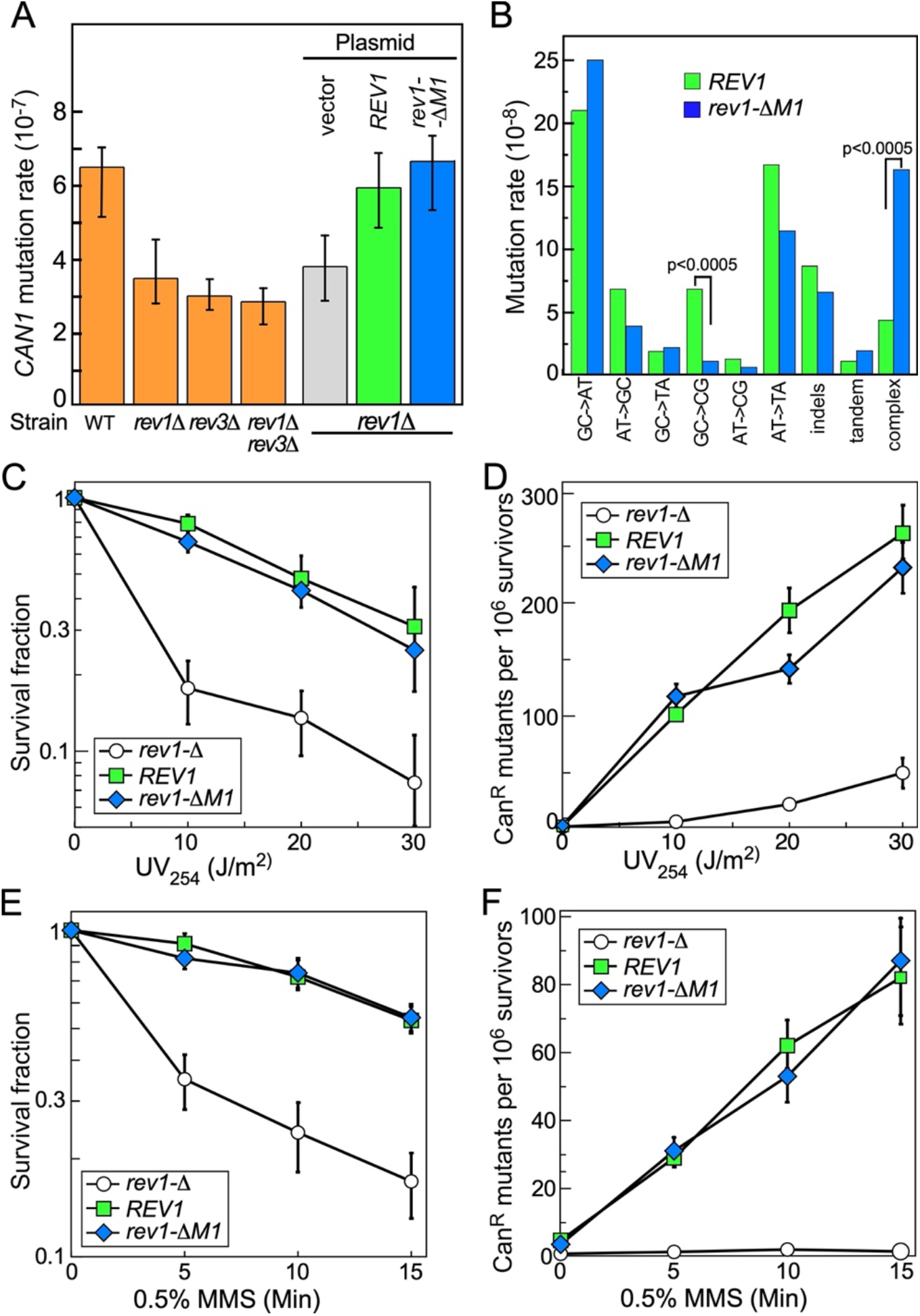
Spontaneous and UV-induced mutations in the *rev1-ΔM1* mutant. (A) Spontaneous mutation rates (with 95% confidence intervals) of wild-type and the indicated *rev3* and/or *rev1* deletion strains in orange, and of the *rev1Δ* strain containing the indicated *REV1* genes as shown. (B) Mutation spectrum in the *CAN1* gene of *REV1* versus *rev1-ΔM1*. Data derived from supplemental tables S3 and S4. (C,E) Survival of the indicated strains after UV-irradiation or MMS treatment. (D,F) Induced mutations of the strains in (C,E). The values in C-F are the mean and standard deviation from two independent replicates.

UV damage sensitivity assays and mutation induction were carried out as described previously (26). For methyl methanesulfonate (MMS) studies, stationary phase cultures were washed with water, resuspended at OD_660_=5 in 50 mM Na-phosphate pH 7.2, and treated with 0.5% of MMS (final) at 30 °C. At 5, 10, 15 min, aliquots were quenched with an equal volume of 10% sodium thiosulfate. After washing, appropriate dilutions were plated as above on minimal complete, selective medium (SC-His-Arg) for viable cell count and on the same medium, but with canavanine for mutant count.

In order to obtain mutation spectra, canavanine plating experiments were repeated for ∼150 total independent cultures. Genomic DNA was prepared from one single colony per canavanine plate and the complete 1.8 kb *CAN1* gene was sequenced. Unambiguous sequence information was obtained from 110 wild-type and 124 ΔM1 isolates, and listed in supplementary Tables S3 and S4, and summarized in Figure 5B. For the statistical significance of the difference between the data in Figure 5B, a student t-test was performed with KaleidaGraph.

## RESULTS AND DISCUSSION

### Rev1 plays a key role in Pol ζ regulation

In a previous study of the bypass synthesis of an interstrand DNA crosslink, we observed that TLS past the G=G crosslink by Pol ζ was stimulated by the addition of Rev1, but that, surprisingly, replication of undamaged DNA by Pol ζ was actually inhibited by addition of Rev1 (50). Here, we study in detail the nature of this opposing regulatory activity of Rev1 and the determinants/domains of Rev1 that are responsible for it. Our studies are aided substantially by the recent cryoEM structure of the Rev1-Pol ζ complex at 3.5 Å resolution (Figure 1A) (22). It depicts interactions between Rev3 and its accessory subunits, Pol31, Pol32, and the two Rev7 subunits, interactions with Rev1, and with template-primer DNA. In this structure, the catalytic domain of Rev1 and its ubiquitin-binding domains (UBM) remain unresolved. The CTD of Rev1 shows interactions with Rev3, with Pol32, and with one Rev7 subunit, as shown previously by domain interaction mapping (24,60). In addition to binding Rev3, the NTD of Rev1 also shows interactions with DNA (61).

Figure 1B presents a primer extension reaction by Pol ζ. A 100-mer circular DNA template was used in order to prevent PCNA from sliding off the DNA after having been loaded by the clamp loader RFC (replication factor C) (62). Single-stranded DNA was coated with RPA (replication protein A), PCNA was loaded (or not), and the assay started by addition of Pol ζ without or with Rev1, either wild-type or the catalytic-dead form (cd = DE467,468AA).

We used Rev1^cd^ to understand which regulatory feature is related to the presence of the Rev1 protein and which to the Rev1 catalytic activity. This assay demonstrates some of Rev1’s regulatory features. On undamaged DNA (lanes 1-12), PCNA strongly stimulated Pol ζ activity (lanes 7,8 versus 1,2), as shown previously (26). In Figure. 1C, the gel data were quantified as a ratio of long extension products (>50-mer) over short extension products (<50-mer). Addition to the assay of wild-type Rev1 inhibited extension without PCNA (lanes 5,6 versus 1,2), and this Rev1-mediated inhibition was even stronger in the presence of PCNA (compare lanes 11,12 with 5,6 and 7,8). Inactivation of Rev1’s catalytic activity (Rev1^cd^) resulted in a less severe inhibition (lanes 9,10 versus 7,8 and versus 11,12), suggesting that the inhibition is composed of two sources, Rev1 protein binding and Rev1-mediated dCMP incorporation. Supplementary Figure S1A shows an enlargement of lanes 7-12 near the primer start position, showing that inclusion of Rev1, but not omission of Rev1 or inclusion of Rev1^cd^, leads to the generation of faster migrating replication products at the +1, +2 and +3 positions, presumably due to dCMP misincorporation. Possibly, stalling after dCMP misincorporation by Rev1 may cause more severe inhibition compared with Rev1 protein binding alone.

Next, we tested the efficiency of Pol ζ to bypass a model abasic site. Whereas we and others have previously shown that Pol ζ has the ability to bypass abasic sites, it is slow in the absence of Rev1 (26,63). In that context, it is noteworthy that Pol ζ replication products show a prolonged stall at the position prior to the abasic site (Figure 1B, lanes13,14,19,20), and so do replication products when Rev1^cd^ is added (lanes 15,16,21,22). However, with Rev1^wt^ temporary stalling occurs opposite the abasic site after rapid dCMP incorporation (lanes 17,18,23,24), but these products are efficiently elongated to longer products. When we measure the percentage of complete abasic site bypass (products >29 nt, Figure 1D), both Rev1^wt^ and Rev1^cd^ show stimulation (lanes 13,14 versus 15-18), and addition of PCNA shows an even stronger stimulation, with the PCNA-Rev1^wt^-Pol ζ complex being most efficient in TLS (lanes 23,24). Interestingly, after bypass of the abasic site by PCNA-Pol ζ, without Rev1, further replication products made by PCNA-Pol ζ are long (Figure 1A, lanes 19,20). In contrast, after bypass of the abasic site by PCNA-Rev1-Pol ζ, further replication products are short (lanes 23,24), indicating that after stimulating damage bypass, Rev1 once again exerts its inhibitory activity on downstream undamaged DNA. Thus, the combined effects of Rev1 and PCNA on the activity of Pol ζ are opposing in nature, when measured on undamaged *versus* damage DNA.

### Domain analysis of Rev1

In a previous study, when mapping the binding sites for K164-ubiquitinated PCNA on Rev1, which is important for efficient mutagenesis and post-replication repair (64), we had designed a series of Rev1 domain mutants (65). We expanded this analysis with additional domain mutants. The Rev1 domain structure and mutants used in the current study are presented in Figure 2A. Upstream of the BRCT domain, we identified a 15-amino-acid conserved motif (position 135-149), named M1 (66). The M1-motif sequence showed conserved amino acids along the alpha-helical structure (22). For our M1 motif studies, we designed two mutants of the M1 motif, Rev1-ΔM1 and Rev1-5A. Rev1-5A was designed having five-point mutation in the most conserved amino acids (Figure 2A). Mutations of the M1 domain do not alter the Rev1 catalytic activity (Figure S6). Additional mutants have a deletion of the BRCT domain (31,35), of a domain containing the UBM motif and UBM-like motifs (65), and of the CTD (34). These mutants, and the catalytic-dead mutant were tested in primer extension (Figure 2B, C, lanes 1-8) and abasic site bypass (lanes 9-16) assays, in order to determine their role in Pol ζ-mediated replication and TLS. All assays in Figure 2 contain PCNA; the corresponding assays without PCNA are presented in Supplementary Figure S2. On undamaged DNA (lanes 1-8), Pol ζ alone extended the primer during the assay, until it reached the end of the template circle, yielding a ∼100 nt extension product. Addition of Rev1^wt^ to Pol ζ resulted in the accumulation of mainly short extension products. This was accompanied by misincorporation of dCMP with Rev1^wt^ and with all the domain mutants, but not with Rev1^cd^ and Pol ζ alone (Figure S1B). The ratio of long to short products was decreased ten-fold when Rev1^wt^ is part of the Pol ζ complex (compare lane 3 with lane 2). In stark contrast to the inhibition exerted by Rev1^wt^, Rev1^ΔM1^ showed almost no accumulation of short products, with the exception of the +1 product resulting from dCMP misinsertion (see supplementary Figure S1B). In addition, longer products were synthesized by Rev1^ΔM1^-Pol ζ than by Pol ζ alone, and some extension products were larger than 100 nt (lane 4), suggesting that Rev1^ΔM1^-Pol ζ has the capability of strand displacement synthesis. Lanes 5-8 represent the reactions of Pol ζ with the ΔBRCT, cd, ΔUBM, and ΔCTD mutants of Rev1, respectively. Only the ΔUBM mutant shows the strong inhibitory activity of Rev1^wt^ (lane 3 versus 7), and Rev1^cd^ shows intermediate inhibitory activity (lane 6), like observed in Figure 1. The ΔBRCT mutant shows a modest increase in long products, but not as strongly as the ΔM1 mutant (compare lane 5 with 4). Most strikingly, the ΔCTD mutant shows very little inhibition (lane 8). As the Rev1 CTD is required for strong binding of Rev1 to Pol ζ, this result suggests that the Rev1-Pol ζ complex be intact for full regulatory activity of Rev1 (67,68).

In the TLS analysis (Figure 2C, lanes 9-16), we quantified both the percentage of AP bypass and the ratio of long/short bypass products. Like observed in Figure 1B, Pol ζ poorly bypassed the abasic site, but those that did bypass were largely extended full length (lane 10). While all forms of Rev1 stimulated bypass synthesis to varying degrees (compare lane 10 with 11-16; upper bar graph), further extension after lesion bypass was analogous to that observed on the undamaged DNA. Thus, while Rev1^wt^ inhibited further extension of the bypass products (compare lane 11 to lane 10), the ΔM1 mutant strongly stimulated long product formation, and the ΔBRCT mutant did so more modestly compared to no Rev1 (lanes 12,13 compared to lane 10). The ΔUBM mutant behaved roughly like wild-type, whereas the ΔCTD mutant showed longer extension products, similar to the no-Rev1 control (compare lane 10 with lane 16). The latter could be due to the formation of an unstable complex between Rev1^ΔCTD^ and Pol ζ.

We also used a replication assay of a 3 kb long template to assess replication efficiency of Pol ζ alone or with (mutant) Rev1 (Figure 2D). In this assay, replication products were only observed if longer than ∼500 nt. These assays largely recapitulated our finding with small oligonucleotide templates: Rev1 inhibited replication by Pol ζ, but two different motif M1 mutants, ΔM1 and 5A, substantially stimulated the synthesis of long DNA products. Replication assays with mutant forms of Rev1 containing combined, multiple mutations were carried out to assess the nature of inhibition or stimulation (Figure 2E). Whereas Rev1^wt^ or Rev1^cd^ almost completely inhibited the formation of long products, the additional deletion of the CTD (ΔC) yielded significantly long product formation (compare lanes 3-6 with 7-10). Conversely, the strong stimulation of long product formation with the Rev1-5A mutant or with the Rev1^cd^-5A double mutant was attenuated when in addition the CTD was deleted (ΔC) (compare lanes 11-14 with 15-18). These data indicate that the stability of the Rev1-Pol ζ complex is important for Rev1 exerting its full either stimulatory or inhibitory activity.

The role of PCNA in Rev1-Pol ζ activity is described in Supplementary Figure S2. In the absence of PCNA, the strong inhibitory effect of Rev1^wt^ on the replication of undamaged DNA (lane 16 versus 15) and the stimulatory effect of the ΔM1 mutant (lane 17 versus 15) are almost entirely attenuated. However, bypass of the abasic site is still stimulated by Rev1^wt^ and by those domain mutants with catalytic activity (lanes 22-28). Supplementary Figure S3 shows that ubiquitination of PCNA does not alter Rev1’s DNA replication and damage bypass properties with Pol ζ and with the various (mutant) Rev1-Pol ζ complexes. These latter results support the proposal that ubiquitination of PCNA is primary important for intracellular signaling and recruitment to stalled forks and to sites of damage (65,69,70).

Whereas tandem BRCT domains are ubiquitous phosphopeptide modules that are involved in the DNA damage response (32,71), the single BRCT domain of Rev1 has been shown to bind PCNA as well as DNA. PCNA binding to BRCT occurs with low affinity (30,35,61). A second PCNA binding domain with the PIP box located on the C-terminal side of the little finger domain, has been identified in the catalytic domain of Rev1 (37,38). This latter domain may have a more important regulatory function during Rev1-Pol ζ replication (see below). The cryoEM structure of yeast Rev1-Pol ζ was resolved together with a bound template-primer and a complementary dNTP, with the primer terminus and the dNTP being localized within the active site of the Pol ζ catalytic subunit Rev3 (22). In this structure the M1-BRCT domain is shown to bind the double stranded region of the template primer (Figure 1A). In particular, several M1 amino acids contact the major groove of dsDNA. Curiously, a DNA binding study of the human Rev1 BRCT region shows strong binding to a single-double stranded DNA junction with a recessed 5’-phosphate (61). The related single BRCT domain of the human Replication factor C Rfc1 subunit likewise binds DNA with a single-double stranded DNA junction containing a recessed 5’-phosphate, and the cryoEM structure of a yeast RFC-DNA complex is consistent with these biochemical data (72,73). Therefore, we investigated the DNA binding properties of the yeast Rev1 NTD (aa 1-251) containing both M1 and BRCT motifs, and of the NTD lacking the M1 motif (Figure S4). Domains were overexpressed and purified from *E. coli* (Figure S4A), and used in an electrophoretic mobility shift assay (EMSA) to assess binding activity to a substrate that was successfully used in the human Rev1-NTD DNA binding study (61).

Both Rev1-NTD and Rev1-NTD^ΔM1^ showed equivalent binding to 5’-phosphorylated and recessed DNA in this EMSA assay (Figure S4B). Furthermore, competition binding assays with either unlabeled substrate DNA or unlabeled substrate DNA lacking the embedded 5’-phosphate showed that DNA containing the 5’-phosphate bound Rev1-NTD with about 5-10 times higher affinity than DNA without 5’-phosphate, and the same was observed with Rev1-NTD^ΔM1^ (Figure S4C,D). From these studies, we conclude that the yeast Rev1-NTD also binds DNA with a recessed 5’phosphate, and that the M1 motif does not contribute to DNA binding. In our DNA replication studies of a 3 kb template, we used a 26 nt 5’-hydroxy primer and ^32^P-dNTPs to label the nascent DNA. When instead we used a 5’-phospho-primer, identical results were obtained (Figure S4E). From these experiments, we conclude that the potential importance of a complex between the BRCT domain of Rev1 and a DNA structure containing a recessed 5’-phosphate DNA has not been revealed in our biochemical studies of the PCNA-Rev1-Pol ζ complex.

### Rev1 reduces Pol ζ processivity

The inhibition of Pol ζ by Rev1 on undamaged DNA may be caused by a decrease in the processivity of DNA replication by Rev1-Pol ζ compared to Pol ζ alone. The processivity analysis was carried out by measuring the distribution of DNA replication products as a function of time at a high DNA/enzyme ratio. Ideally, if primer utilization increases with time while the product distribution does not, then processivity conditions are met, and the processivity can be defined as the median nucleotide of the product distribution (i.e. products shorter and longer than the median are equal in magnitude). In practice, the distribution of replication products may increase with time, and the processivity needs to be obtained by extrapolation to minimal times (Figure 4).

One of the issues we needed to address is what happens to the complex during our processivity analysis, which is carried out in the presence of a five-fold excess of primer-template DNA? Is it the case that the complex dissociates from the DNA after a processive cycle, and then rebinds either to a new, unreplicated DNA substrate or to an already extended DNA with equal chance, depending on their respective concentrations? Or does the bound complex pause for extended time followed by continuation of replication of the DNA to which it initially was bound? In the latter case, maximum primer utilization would be 20%. Figure 3B shows that the percentage of primer utilization increased with time for each reaction, up to 70% utilization, indicating that a processive cycle is terminated by dissociation of the complex.

To analyze the replication data, the lanes were quantified and the median nucleotide determined from the extension product distribution and plotted as a function of reaction time (Figure 3C). The number obtained by extrapolation to zero defines the processivity. In this particular analysis, PCNA-Pol ζ replicates about 10 nucleotides before dissociation. Processivity of the PCNA-Rev1^wt^-Pol ζ complex is drastically reduced to 3-4 nt, whereas processivity of the PCNA-Rev1^ΔM1^-Pol ζ complex is increased to 28 nt. A second experiment with these three enzyme complexes, carried out at slightly different enzyme concentrations, yielded similar processivity numbers (Figure S5, A-C). Thus, the inhibitory effect of Rev1^wt^ on Pol ζ and the stimulatory effect of Rev1^ΔM1^ can, to a large degree, be ascribed to the relative processivities of these enzyme complexes.

### Rev1 stimulates bypass of DNA polymerase stall sites caused by dNTP omission

We want to understand whether the mechanism of Rev1-Pol ζ stalling and bypass at sites of DNA damage is similar to the occurrence of stalling on undamaged DNA that is caused by an obstacle to DNA synthesis. Therefore, we carried out a replication reaction of in which one of the 4 dNTPs is missing, thereby causing stalling at complementary template positions (Figure 4). We used PCNA-Pol ζ without Rev1 or with Rev1^wt^ or Rev1^cd^, and either 4 or 3 dNTPs at 10 μM each for 10 minutes. The template sequence is shown on either side of the gel. Red asterisks on the gel represent positions that are located one nucleotide upstream of the template position corresponding to the omitted complementary dNTP. Indeed, at each of these starred positions, replication stalls because the next base-pairing dNTP is missing. We made several remarkable observations in this experiment. First, the PCNA-Pol ζ complex shows very poor bypass synthesis. For instance, in the absence of dCTP, all products stalled at the +1 and +2 positions, upstream of template Gs at the +2 and +3 positions (lane 13), whereas with Rev1 added, the PCNA-Rev1^wt^-Pol ζ complex bypassed up to 10 G-template positions (lane 14). Second, results with the catalytic-dead mutant were very comparable to wild-type. Perhaps this was to be expected for the -dCTP assay because Rev1 solely uses dCTP as substrate (lanes 14, 15), but it was also the case with the other omission assays (lanes 5 and 6, 8 and 9, 11 and 12). Therefore, both Rev1^wt^ and Rev1^cd^ strongly stimulate Pol ζ to misincorporate dNTPs, and the extended misincorporation tracks for each of the four omissions suggest that no specific dNTP is required for it. Third, the reading frame was strongly maintained, suggesting that bypass is not mediated by frameshifting. Our results are consistent with genetic studies in chicken DT40 showing that Rev1 suppresses frame-shift mutagenesis by Pol ζ (74). Fourth, Rev1^ΔM1^ stimulated misincorporation by Pol ζ moderately more efficient than Rev1^wt^ (Figure S6). The conclusion is that, while Rev1 functions as an inserter enzyme and Pol ζ as an extender polymerase when actual DNA damage is present, such as an abasic site or a DNA interstrand crosslink (Figure 1B) (17,50), Rev1 functions as a stimulator of Pol ζ misincorporation activity once the complex has paused on undamaged DNA.

### Spontaneous and damage-induced mutagenesis of the rev1-ΔM1 mutant

Our biochemical studies of Pol ζ and (mutant) Rev1 show the effect of Rev1 on Pol ζ’s enzymatic activity on undamaged and damaged DNA. From the biochemical properties of the Rev1^ΔM1^ mutant protein, i.e. increased processivity of DNA synthesis, one might expect that in yeast this mutation may lead to an increase in spontaneous mutation rates and an increase in damage-induced mutagenesis. We used resistance to the arginine analog canavanine by mutational inactivation of the *CAN1* gene to obtain quantitative values for mutation rates. All yeast strains in this study are isogenic and were grown in complete minimal media prior to plating on canavanine-containing complete minimal media without or with UV irradiation. In accord with many previous studies (46,75,76), the spontaneous mutation frequencies of *rev1Δ, rev3Δ* and the *rev1Δ rev3Δ* double mutant were about 40-50% of wild-type (Figure 5A), indicating that about 50-60% of the mutations in a wild-type yeast strain result from the action of Rev1-Pol ζ at forks that are stalled because of spontaneous damage or other impediments to elongation. *REV1* or *rev1-ΔM1* were introduced into the *rev1Δ* mutant strain and both spontaneous, UV-induced, and MMS-induced mutations were measured (Figure 5A,B,D,F). Interestingly, the spontaneous mutation rate of *rev1-ΔM1* was not significantly different from wild-type. We sequenced 110 independently arrived spontaneous *CAN1* mutants for *REV1* and 124 for *rev1-ΔM1* and compared the mutation spectra (Figure 5B, Supplementary Tables S3, S4). Two differences between *REV1* and *rev1-ΔM1* were detected with high significance. First, the rate of GC→CG mutations were substantially reduced in *rev1-ΔM1*. The GC→CG mutations are a hallmark signature for the Rev1 contribution to Rev1-Pol ζ mutagenesis, through incorporation of dCMP opposite a template C (40). The majority of GC→CG mutations in wild-type *REV1* occur in a single hotspot at nt 374 of the *CAN1* gene (Supplementary Table S3). However, *rev1-ΔM1* yielded no mutations at that position, for which we can currently offer no explanation. Second, the frequency of complex mutations is elevated almost four-fold in the *rev1-ΔM1* mutant, in keeping with the increased processivity of the PCNA-Rev1ΔM1-Pol ζ enzyme complex. However, complex mutations are mostly limited to a tract length of 3-4 (Table S4). Only 2 out of the 124 spontaneous *CAN1* mutations isolated from the *rev1-ΔM1* strain have a longer track length than 3-4, of 29 and 34 nt. If one assumes that these longer track lengths did not arise from independent mutational events, it can be concluded that the TLS patch in the *ΔM1* mutant, while longer than in wt *REV1*, still remained fairly small. In addition, we did not measure a significant difference between *REV1* and *rev1-ΔM1* in damage sensitivity (Figure 5C,E), and in damage induced mutagenesis (Figure 5D,F), suggesting that other redundant mechanism(s) than the M1 motif of Rev1 may restrain the extent of error-prone replication with associated mutagenesis by Rev1-Pol ζ.

For many years, Rev1 has been considered as a scaffold protein important for regulating TLS and mutagenesis by other DNA polymerases including Pol ζ (46,49,77). However, most studies, including some from our own laboratory, have focused on delineating the properties of Rev1 and Pol ζ in isolation. Our current findings clearly demonstrate that the activities of the trimeric PCNA-Rev1-Pol ζ complex cannot be deduced from a simple composite of their individual properties. PCNA is essential for Rev1 regulation of both the stimulation and inhibition of Pol ζ activity. Two PCNA-binding elements have been identified in Rev1, the first in the BRCT domain and the second in the PAD (polymerase-associated domain), also called little finger domain of the catalytic module (30,37). Since deletion of the BRCT domain maintained the PCNA regulatory function of Rev1 in DNA damage bypass (Figure 2B, lane 14), it is likely that the PCNA-binding domain in PAD is responsible for that regulation.

As a physiological argument, it makes perfect sense that Rev1 stimulates damage bypass by Pol ζ but inhibits further DNA synthesis once the damage has been bypassed, because this model limits mutations to those that are necessary to bypass the damage, but not beyond. However, because mutation of the M1 regulatory motif does not show a strong mutator phenotype, a likely conclusion is that other, redundant mechanism(s), perhaps that of PCNA de-ubiquitination, maintain the inhibitory post-bypass phenotype.

## DATA AVAILABILITY

All relevant data are included in the paper and the supplementary data section, and none have been deposited in online databases. Plasmids and yeast strains are available upon request.

## SUPPLEMENTARY DATA

Supplementary Data are available at NAR online.

## ACKNOWLEDGEMENTS

We thank members and former members of the Burgers lab for valuable discussions on the project and on the manuscript.

