## Supplemental data, tables and figures for "Opposing regulation by Rev1 of DNA polymerase zeta activity on damaged versus undamaged DNA"

### Table S1. Rev1 Mutants

| Rev1 variant | mutation description |
|---|---|
| Rev1-cd (catalytic dead) | DE467,468AA |
| Rev1-ΔM1 | Δ130-149 |
| Rev1-5A | YF135,136AA, K139A, Q146A, Q148A |
| Rev1-ΔBRCT | Δ160-250 |
| Rev1-ΔUBM | Δ743-840 |
| Rev1-ΔCTD | Δ865-985 – C terminal truncation |
| Rev1-cd,ΔCTD | Combinations of above mutations |
| Rev1-ΔM1,cd | |
| Rev1-ΔM1,cd,ΔCTD | |
| Rev1-5A,cd | |
| Rev1-5A,cd,ΔCTD | |

### Table S2. Oligonucleotide sequences

| Oligo name | Sequence |
|---|---|
| SKII 100 | 5'/Phos/AACAAAAGCTGGAGCTCCACCGCGGTGGCGGCCGCTCTAGAACTAGTG GATCCCCCGGGCTGCAGGAATTCGATATCAAGCTTATCGATACCGTCGACCT-3' |
| SKII 99 AP1 | 5'/Phos/AACAAAAGCTGGAGCTCCACCGCGGTGGCGGCCGCTCTAGAACTAGTG GATCCCCCGGGCT/idSp/CAGGAATTCGATATCAAGCTTATCGATACCGTCGACCT-3' |
| SKII 100 Anneal | 5'-GCTCCAGCTTTTGTTAGGTCGACGGTATCG -3' |
| SKII primer 1 | 5'-TATCGATAAGCTTGATATCGAATTCC -3' |
| SKII primer 2 | 5'-GGTATCGATAAGCTTGATATCGAATT-3' |
| SKII primer 3 | 5'-CGGTATCGATAAGCTTGATATCGAAT-3' |
| SKII primer 4 | 5'-AGGTCGACGGTATCGATAAGCTTGAT-3' |

The /idSp/ designation by IDT = tetrahydrofuran, functioning as a model abasic site.

### Supplementary proteins

#### Split Ubi PCNA and Split PCNA purification

Ub-PCNA was designed as two polypeptides (Split Ub-PCNA ) as described in (1) and purified as described in (2) with modifications. Split Ub-PCNA construct was transformed into *E. coli* BL21-DE3+RIL and grown on 1 L of LB + Carbenicillin + Chloramphenicol (working concentration used as recommended by the manufacturer) at 37°C to $A_{600}$ = 0.8, followed by addition of 1 mM IPTG and incubation for an additional 4 hours. Cells were harvested by centrifugation and resuspended in Buffer A (25 mM Hepes 7.8, 40 mM Triethanolamine, 1 mM EDTA, 10% Glycerol, 50 mM NaCl, 1 mM 2-Mercaptoethanol, 2 μM E-64, 5 μM Bestatin, 10 μM Pepstatin A, 10 μM Leupeptin and 1 mM PMSF) with 1 mg/ml lysozyme. The lysate was further processed through ammonium sulfate precipitation as described in (2). The ammonium sulfate pellet was resuspended in Buffer A + 5 mM Imidazole. The sample was loaded on Ni-NTA column and washed several times with Buffer A + 5 mM imidazole and increasing concentration of Imidazole (5 mM,10 mM,100 mM). The sample was eluted with Buffer A + 200 mM Imidazole. 1 mM EDTA was added back to the sample and loaded on 1 ml MonoQ column (Cytiva). Split Ub-PCNA was eluted with 20 ml of linear gradient of 50 mM-500 mM NaCl in Buffer A.

#### Rev1-NTD expression and purification

Rev1 N-Terminal domain 1-251 aa (NTD) fragment was cloned into pGEX-6P1 vector for expression as an N-terminal GST fusion protein. (pGEX-6P-1 tacP-lacO-GST-PreScission-Rev1(1–251) AmpR ori(pBR322)). We cloned the wild-type and the ΔM1 forms. The construct was transformed into *E.coli* BL21-DE3 and grown on 1 L of LB + carbenicillin (working concentration used as recommended by the manufacturer) at 37°C to $A_{600}$ = 0.6-0.8, followed by the addition of 1 mM IPTG and incubation for an additional 4 hours. Cells were harvested by centrifugation and resuspended in Buffer A (50 mM Hepes 7.8, 8% glycerol, 1 mM DTT, 0.2 M NaCl, 2 μM E-64, 5 μM Bestatin, 10 μM Pepstatin A, 10 μM Leupeptin and 1 mM PMSF) with 1 mg/ml lysozyme. The lysate was incubated on ice for 30min, then 0.05% Tween and 0.01% $C_{12}E_{10}$ were added, followed by sonication. After sonication, the lysate was further processed through ammonium sulfate precipitation. The ammonium sulfate pellet was resuspended in Buffer B (50 mM Hepes 7.8, 8% glycerol, 1 mM DTT, 1 mM EDTA, 0.05% Tween, 0.01% $C_{12}E_{10}$, 2 μM E-64, 5 μM Bestatin, 10 μM Pepstatin A, 10 μM Leupeptin and 1 mM PMSF). The sample was incubated with GST beads (Cytiva) for 2 hours. The beads were washed extensively with Buffer B + 0.5 M NaCl, followed by a wash with Buffer B + 0.2 M NaCl. The protein was eluted with Elution Buffer (Buffer B + 0.2 M NaCl + reduced Glutathione). The eluted sample was incubated with 3C-PreScission protease for 16 hours. The sample was loaded on 1 ml MonoS (Cytiva) and was eluted with 20 ml of a linear gradient of 50 mM-500 mM NaCl in 50 mM Hepes 8.0, 8% glycerol, 1 mM DTT, 1 mM EDTA, 0.01% $C_{12}E_{10}$, 2 $\mu$M E-64, 5 $\mu$M Bestatin, 10 $\mu$M Pepstatin A, 10 $\mu$M Leupeptin and 1 mM PMSF.

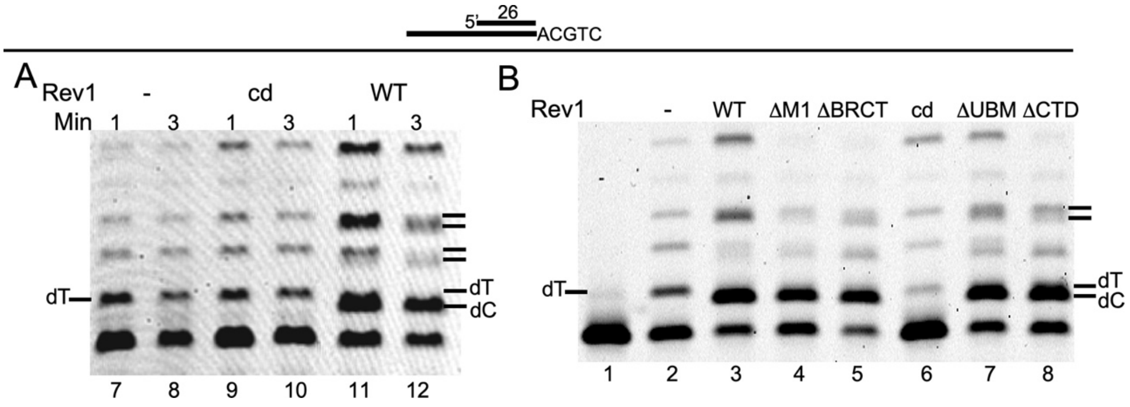


**Figure S1.** Rev1-mediated misincorporation of dCMP. Enlargement and focus on the initial few nucleotides replicated in the assay described in Figure 1B and Figure 2B. DNA substrate used in the assay was SKII-100 circular ssDNA primed with SKII primer 1(A) Lanes 11,12 from Figure 1B show the misincorporation of dCMP (as faster migrating bands) by $Rev1^{wt}$-Pol $\zeta$, in comparison to Pol $\zeta$ alone (lanes 7,8) or $Rev1^{cd}$-Pol $\zeta$ (lanes 9,10). (B) Similarly, an enlargement from Figure 2B. Inclusion of Rev1 in the assay (lane 3), and of all Rev1 mutants except $Rev1^{cd}$ (lane 6) show faster migrating bands due to dCMP misincorporation.

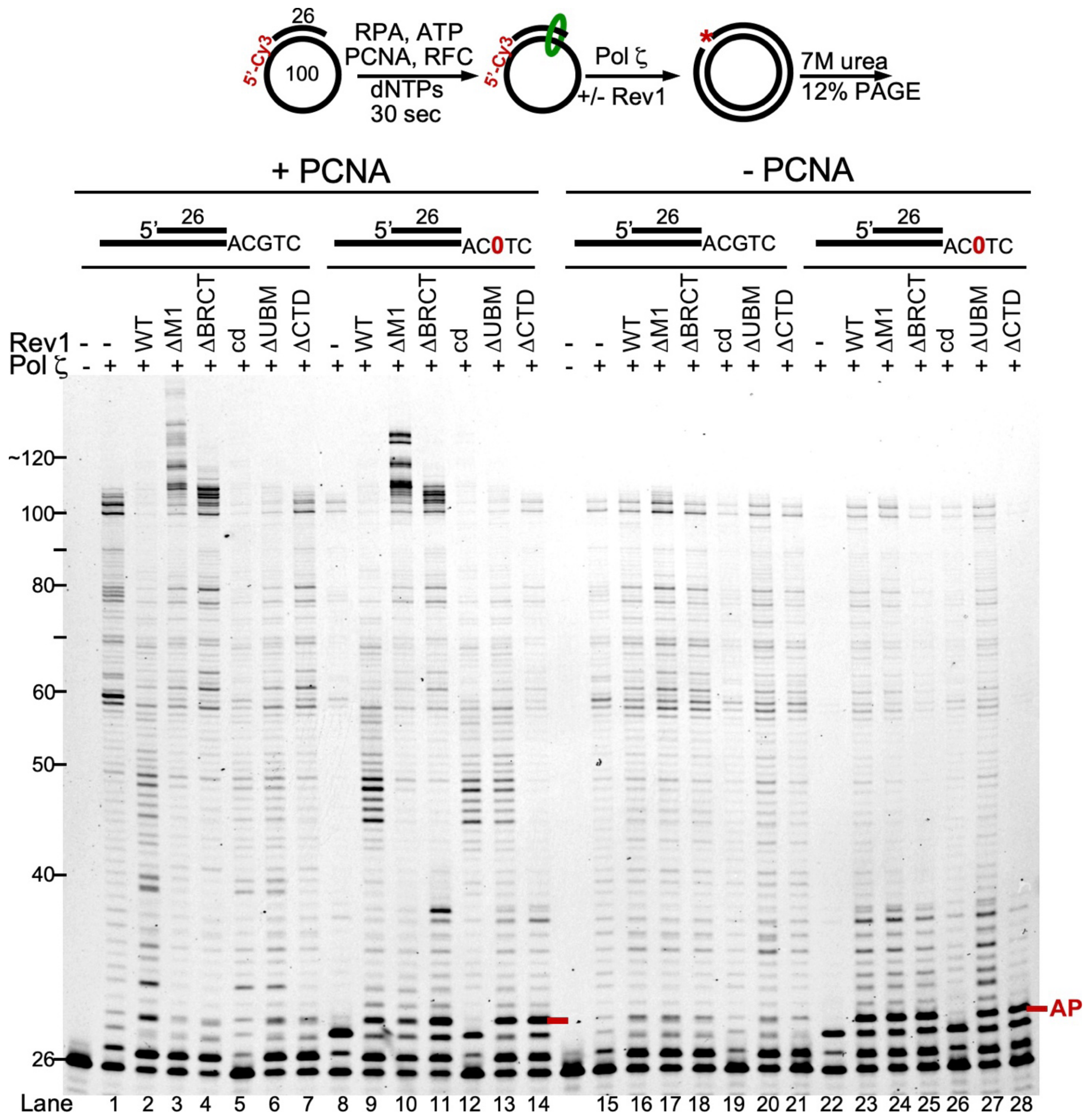


**Figure S2.** DNA synthesis with mutant forms of Rev1. The left half of this gel (+PCNA data) is presented in Figure 2B (see legend to that Figure). Remarkably, without PCNA, Pol ζ alone, Rev1$^{wt}$-Pol ζ, and the different Rev1$^{mutant}$-Pol ζ complexes (lanes 15-21) replicate the 100mer template with comparable efficiency. These results are starkly different when PCNA is present (lanes 1-7). TLS bypass efficiency of the abasic site is low with Pol ζ alone, with Rev1$^{cd}$-Pol ζ, and with Rev1$^{\Delta CTD}$-Pol ζ (lanes 22, 26, 28), whereas bypass efficiency is similar for Rev1$^{wt}$ and the other 3 mutants (lanes 23-25, 27). Again, this contrasts strongly with TLS in the presence of PCNA (lanes 8-14). DNA substrates used in the assay were SKII-100/SKII-AP1 circular ssDNA primed with 5'-Cy3-SKII primer 1.

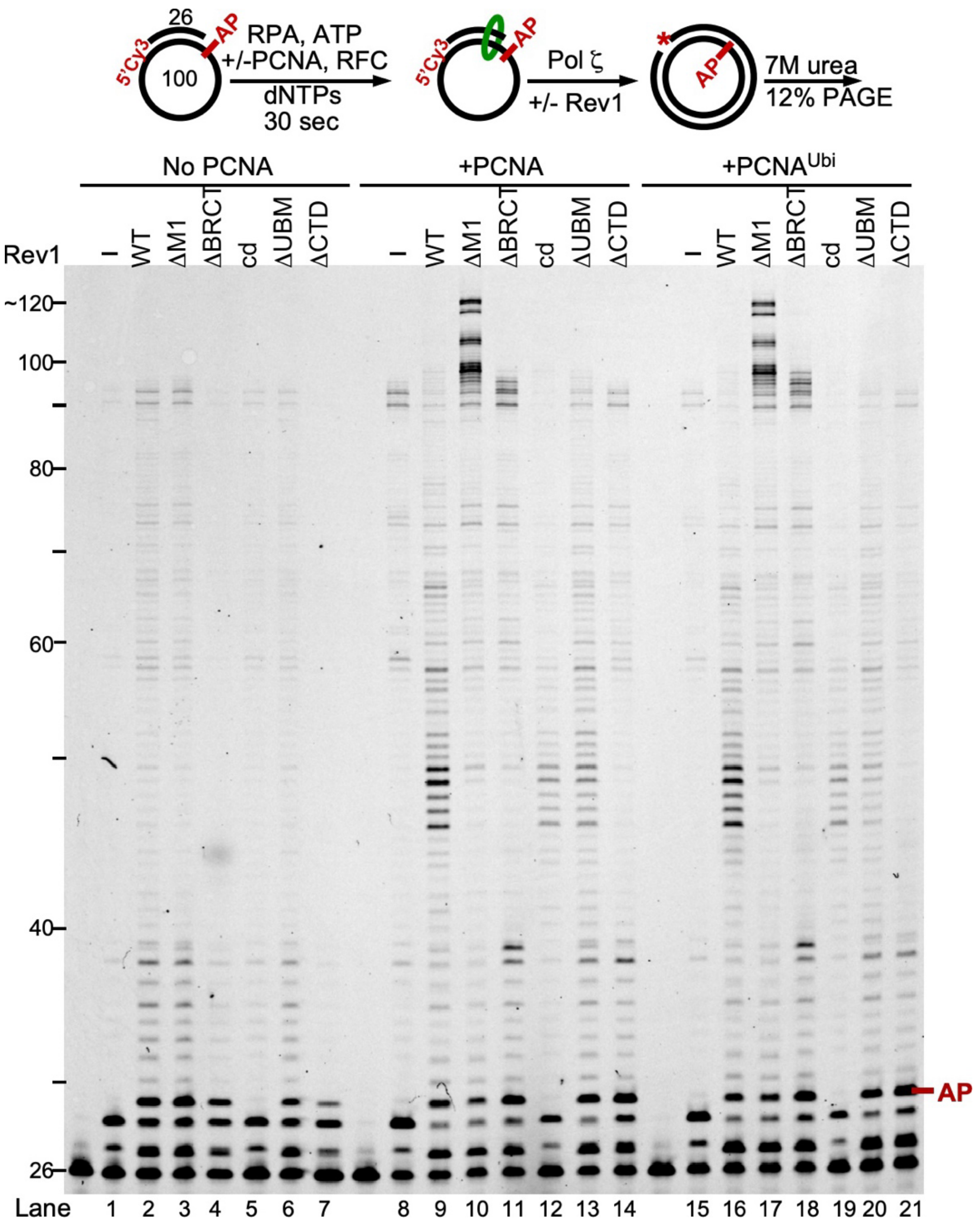


**Figure S3.** Replication of an abasic site with PCNA and ubiquitinated PCNA. 12% urea- PAGE of an abasic site (DNA substrate used in the assay was SKII-AP1 circular ssDNA primed with 5'-Cy3-SKII primer 1) bypass assay with Pol ζ -/+ Rev1 variants using three different conditions: no PCNA, unmodified PCNA, and mono-ubiquitinated PCNA ($PCNA^{Ubi}$). $PCNA^{Ubi}$ was designed as two polypeptides (1-163 + Ub-K164-258) (Split Ub-PCNA ) as described in (1) and essentially purified as described in (2). See description above under Supplementary proteins. The -PCNA and +PCNA data are similar to shown in Figure S2. The salient part of this experiment is that both PCNA and $PCNA^{Ubi}$ show identical replication and damage bypass properties with the various Rev1-Pol ζ complexes (compare lanes 8-14 with 15-21).

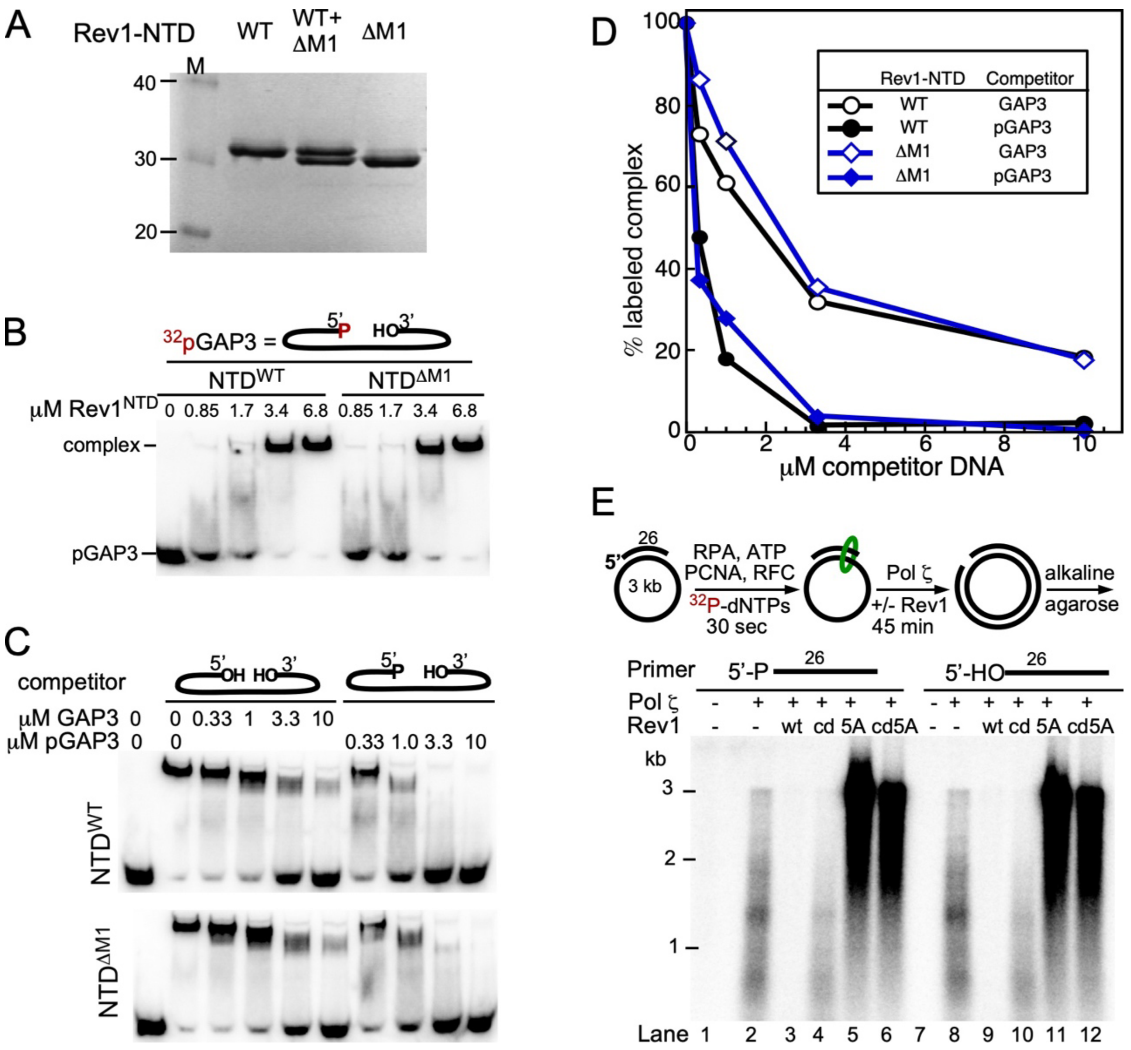


**Figure S4.** DNA binding analysis of the Rev1 NTD and the role of a 5'-phospho primer in replication. (A-D) EMSA analysis of the Rev1 NTD (aa 1-251). (A) SDS PAGE analysis of the Rev1$^{wt}$-NTD and Rev1$^{\Delta M1}$-NTD. About 500 ng of each protein and 500 ng of a 1:1 mixture was analyzed by 12% SDS-PAGE. (B) The DNA substrate used for EMSA DNA binding analysis was a double hairpin with a 3 nt ssDNA gap (GAP3), taken from (3). GAP3 DNA was 5'-labeled with $^{32}$P by T4 polynucleotide kinase and [$\gamma$-$^{32}$P]-ATP, yielding $^{32}$pGAP3. EMSA was carried out as described (3), with 4 nM $^{32}$pGAP3 and indicated concentrations of the NTDs. (C) Competition analysis. The $^{32}$pGAP3-NTD$^{wt}$ (upper gel) or $^{32}$pGAP3-NTD$^{\Delta M1}$ (lower gel) complex was challenged with increasing concentrations or either cold GAP3 or pGAP3. (D) Quantification of the competition analysis. This analysis shows that pGAP3 is a 5-10 fold better competitor than GAP3 and in addition, that there is no difference in binding between Rev1-NTD and Rev1-NTD$^{\Delta M1}$. (E) Top, schematic of the assay. The primer was unlabeled, and the products were metabolically labeled with $^{32}$P-dNTPs. Assays were carried out for 45 min and the products were separated on an 1% alkaline agarose gel.

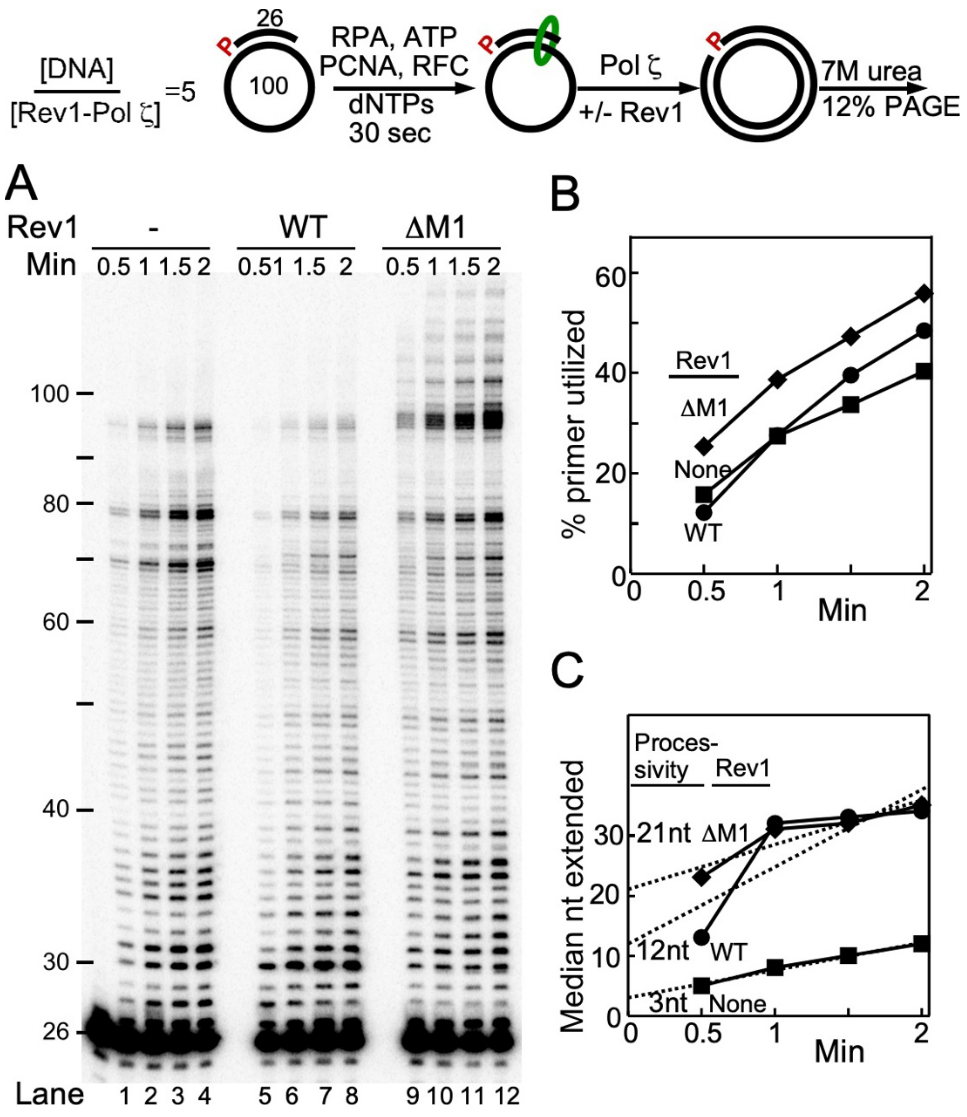


**Figure S5.** Rev1 determines Pol ζ replication processivity. This assay and its analysis are identical to the one described in Figure 4, except that we used 2nM Pol ζ, or Rev1$^{wt}$-Pol ζ or Rev1$^{\Delta M1}$-Pol ζ, and 10nM of the DNA substrate (DNA substrate used in the assay was SKII-100 circular ssDNA primed with 5'-$^{32}$P-SKII primer 4). (A) 12% UREA-PAGE analysis of a time-course replication assay. (B) Plot of the % primer utilized with time. (C) The median nucleotide positions, of extended products, was calculated from the intensity traces of each time point, and plotted against reaction time. Linear plots were extrapolated to time zero to determine processivity.

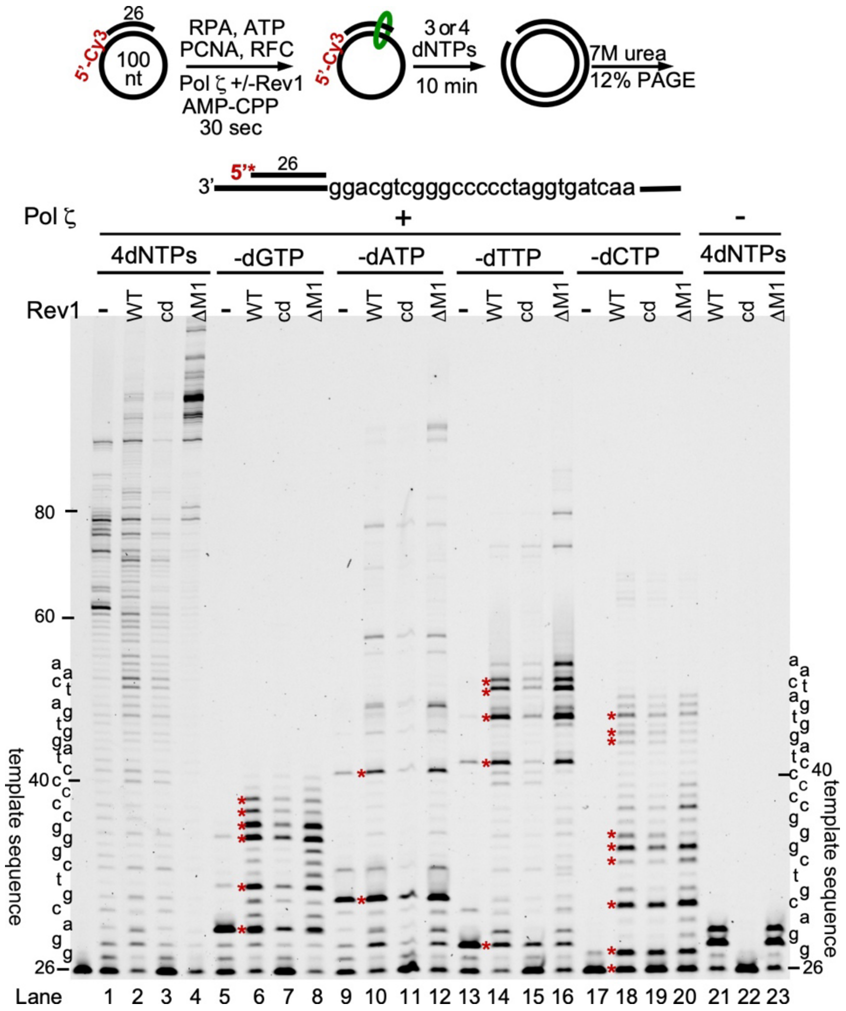


**Figure S6.** Nucleotide omission assay. Top, reaction scheme. Note that this assay has a different primer than the assay shown in Figure 4. (DNA substrate used in the assay was SKII-100 circular ssDNA primed with 5'-Cy3-SKII primer 2). The primer starts with a dG-dG template sequence, allowing for a more robust measurement of Rev1 only activity (lanes 21-23), without Pol ζ. The 12% urea-PAGE analysis shows Pol ζ alone, or in a complex with Rev1$^{wt}$, Rev1$^{cd}$, and Rev1$^{\Delta M1}$, respectively, using either the 4 dNTPs, or lacking dGTP, dATP, dTTP, or dCTP, respectively, as shown. Lanes 21-23 show assays with all 4 dNTPs and with Rev1$^{wt}$, Rev1$^{cd}$, or Rev1$^{DM1}$, but lacking Pol ζ. The template sequence is shown beside the gel. The red stars represent stall positions; each of these positions is located one nucleotide before the template nucleotide, which is supposed to be paired with the missing dNTP in the assay. Note that Rev1$^{wt}$ and Rev1$^{\Delta M1}$ have the same activity (lanes 21,23), whereas Rev1$^{cd}$ is inactive (lane 22). PCNA was present in all assays.

**Table S3.** Spontaneous single nucleotide mutations in the *CAN1* gene. The sequence of the *CAN1* gene is shown, base numbers indicated to the right. Base changes are shown below the sequence: single mutations isolated in wild-type *REV1* (a total of 102) are shown in black, and those isolated in *rev1-ΔM1* (a total of 92) are shown in red. Single nt deletions are shown by a dash and insertions are underlined.

```
ATGACAAATT  CAAAAGAAGA  CGCCGACATA  GAGGAGAAGC  ATATGTACAA  50
                t                                      -

TGAGCCGGTC  ACAACCCTCT  TTCACGACGT  TGAAGCTTCA  CAAACACACC  100
ACAGACGTGG  GTCAATACCA  TTGAAAGATG  AGAAAAGTAA  AGAATTGTAT  150
                   -                              t

CCATTGCGCT  CTTTCCCGAC  GAGAGTAAAT  GGCGAGGATA  CGTTCTCTAT  200
GGAGGATGGC  ATAGGTGATG  AAGATGAAGG  AGAAGTACAG  AACGCTGAAG  250
                 -              -                     t

TGAAGAGAGA  GCTTAAGCAA  AGACATATTG  GTATGATTGC  CCTTGGTGGT  300
              c t          t
              c            t
                           tt

ACTATTGGTA  CAGGTCTTTT  CATTGGTTTA  TCCACACCTC  TGACCAACGC  350
      t  t  t  a                a

CGGCCCAGTG  GGCGCTCTTA  TATCATATTT  ATTTATGGGT  TCTTTGGCAT  400
     t                     g   a
                           g   g           c
                         ggggg

ATTCTGTCAC  GCAGTCCTTG  GGTGAAATGG  CTACATTCAT  CCCTGTTACA  450
                    a                            t

TCCTCTTTCA  CAGTTTTCTC  ACAAAGATTC  CTTTCTCCAG  CATTTGGTGC  500
 t  t           -               t
 t                              t
 t

GGCCAATGGT  TACATGTATT  GGTTTTCTTG  GGCAATCACT  TTTGCCCTGG  550
                    ac     t  tac          tt
                              t c                        c
                              t
                           tttttt

AACTTAGTGT  AGTTGGCCAA  GTCATTCAAT  TTTGGACGTA  CAAAGTTCCA  600
  t                t          t         a

CTGGCGGCAT  GGATTAGTAT  TTTTTGGGTA  ATTATCACAA  TAATGAACTT  650
             a     a         -a                     a  a
             a               aa
GTTCCCTGTC  AAATATTACG  GTGAATTCGA  GTTCTGGGTC  GCTTCCATCA  700
```

```
     t t         c      a a         c    a
     t

AAGTTTTAGC  CATTATCGGG  TTTCTAATAT  ACTGTTTTTG  TATGGTTTGT  750
      a         a            t
      a         t
      a

GGTGCTGGGG  TTACCGGCCC  AGTTGGATTC  CGTTATTGGA  GAAACCCAGG  800
         -                                c a

TGCCTGGGGT  CCAGGTATAA  TATCTAAGGA  TAAAAACGAA  GGGAGGTTCT  850
      a                      t                          -
                             t
                             t

TAGGTTGGGT  TTCCTCTTTG  ATTAACGCTG  CCTTCACATT  TCAAGGTACT  900
                                 c     -             ac

GAACTAGTTG  GTATCACTGC  TGGTGAAGCT  GCAAACCCCA  GAAAATCCGT  950
a   g                    at               tt       -
                           at             tg

TCCAAGAGCC  ATCAAAAAAG  TTGTTTTCCG  TATCTTAACC  TTCTACATTG  1000
        a         t             ta               a-cg
                                t                 t gg
                                tt

GCTCTCTATT  ATTCATTGGA  CTTTTAGTTC  CATACAATGA  CCCTAAACTA  1050
   t ta            c        at-     t               t
                   a        at      t               t
                  aaa       -

ACACAATCTA  CTTCCTACGT  TTCTACTTCT  CCCTTTATTA  TTGCTATTGA  1100
GAACTCTGGT  ACAAAGGTTT  TGCCACATAT  CTTCAACGCT  GTTATCTTAA  1150
CAACCATTAT  TTCTGCCGCA  AATTCAAATA  TTTACGTTGG  TTCCCGTATT  1200
              tca               t  a c           tt   t
                a                                t

TTATTTGGTC  TATCAAAGAA  CAAGTTGGCT  CCTAAATTCC  TGTCAAGGAC  1250
 a  c  a    c                          t
            c                          t
            c                          tt

CACCAAAGGT  GGTGTTCCAT  ACATTGCAGT  TTTCGTTACT  GCTGCATTTG  1300
   -

GCGCTTTGGC  TTACATGGAG  ACATCTACTG  GTGGTGACAA  AGTTTTCGAA  1350
                                            t

TGGCTATTAA  ATATCACTGG  TGTTGCAGGC  TTTTTTGCAT  GGTTATTTAT  1400
```

```
       a                 t       -     -         a

CTCAATCTCG  CACATCAGAT  TTATGCAAGC  TTTGAAATAC  CGTGGCATCT 1450
-                                   c a

CTCGTGACGA  GTTACCATTT  AAAGCTAAAT  TAATGCCCGG  CTTGGCTTAT 1500
              a               t     a t
                              t     g
                              tt   aa

TATGCGGCCA  CATTTATGAC  GATCATTATC  ATTATTCAAG  GTTTCACGGC 1550
                                          t
                                          t

TTTTGCACCA  AAATTCAATG  GTGTTAGCTT  TGCTGCCGCC  TATATCTCTA 1600
TTTTCCTGTT  CTTAGCTGTT  TGGATCTTAT  TTCAATGCAT  ATTCAGATGC 1650
                         a      a     t
                          a           t
                        aaaa          t

AGATTTATTT  GGAAGATTGG  AGATGTCGAC  ATCGATTCCG  ATAGAAGAGA 1700
    tt tt

CATTGAGGCA  ATTGTATGGG  AAGATCATGA  ACCAAAGACT  TTTTGGGACA 1750
AATTTTGGAA  TGTTGTAGCA  TAG                               1773
```

**Table S4.** Tandem and complex mutations in the *CAN1* gene. The first column shows the base position of the initial mutation. The DNA sequences are shown as codons, with the wild-type sequence on the left and the mutations on the right. The bases in bold highlight the changes. The length of the mutation tract is given in parentheses on the right. Tandem mutations have a tract length of (2) and complex mutations have a track length of >(2)

Tandem/Complex mutations (8 total) isolated in wild-type *REV1* strain

| | | | | |
|---|---|---|---|---|
| 519 | ATG TA**T** T**G**G | → | ATG TA**a** T**t**G | (3) |
| 527 | TTT T**C**T TGG | → | TTT**t** T**t**T TGG | (3) |
| 763 | GTT **AC**C GGC | → | GTT **ttt**C GGC | (3) |
| 830 | AAG G**AT** AAA | → | AAG G**g-** AAA | (2) |
| 970 | **G**TT **G**TT TTC | → | **at**TT **t**TT TT**t**C | (10) |
| 1198 | CGT **A**TT TT**A** | → | CGT **t**TT TT**g** | (6) |
| 1406 | TCA A**TC** T**C**G | → | TCA A**-a** T**t**G | (4) |
| 1657 | TTT **A**TT TGG | → | TTT **ttt**TT TGG | (3) |

Tandem/Complex mutations (32 total) in the *rev1*-Δ*M1* strain

| | | | | |
|---|---|---|---|---|
| 305 | ACT A**T**T **G**GT | → | ACT A**a**T **t**GT | (3) |
| 387 | AT**G** **G**GT TCT | → | AT**t** **-**GT TCT | (2) |
| 440 | TTC A**TC** **C**CT | → | TTC A**ca** **a**CT | (3) |
| 450 | GTT AC**A** T**C**C | → | GTT AC**c** T**t**C | (3) |
| 550 | CTG **G**A**A** CTT | → | CTG **a**A**c** CTT | (3) |
| 553 | GAA **C**T**T** AGT | → | GAA **-**T**a** AGT | (3) |
| 600 | GTT CC**A** **C**TG | → | GTT CC**-** **T**TG | (2) |
| 616 | ATT **A**GT ATT | → | ATT **t**GT ATT | (61) |
| +676 | GAA **T**TC GAG | → | GAA **-**TC GAG | |
| 619 | **AT**T TTT TGG GTA | → | **ta**T TTT TGG **G**GTA | (38) |
| +656 | TTC C**C**T GTC | → | TTC C**t**T GTC | |
| 654 | TTG T**TC** CCT | → | TTG T**-a** CCT | (2) |
| 665 | AAA T**A**T **T**AC | → | AAA T**t**T **c**AC | (3) |
| 665 | AAA T**A**T **T**AC | → | AAA T**t**T **c**AC | (3) |
| 715 | ATT **A**T**C** GGG | → | ATT **-**T**a** GGG | (3) |
| 726 | TTT CT**A** ATA | → | TTT CT**t** ATA | (10) |
| +735 | TGT T**T**T TGT | → | TGT T**-**T TGT | |
| 859 | **G**TT TCC **T**C**T** | → | **t**TT TCC **c**C**-** | (10) |
| 893 | TTT C**A**A **G**GT | → | TTT C**g**A **a**GT | (3) |
| 925 | GGT **G**A**A** GCT | → | GGT **a**A**g** GCT | (3) |
| 925 | GGT **G**A**A** GCT | → | GGT **a**A**g** GCT | (3) |
| 950 | TCC G**T**T **C**CA | → | TCC G**g**T **t**CA | (3) |
| 950 | TCC G**T**T **C**CA | → | TCC G**g**T **t**CA | (3) |
| 1019 | ATT G**G**A **C**TT | → | ATT G**a**A **–**TT | (3) |
| 1022 | GGA C**T**T **T**TA | → | GGA C**c**T **c**TA | (3) |
| 1042 | GAC **CC**T AAA | → | GAC **-t**T AAA | (2) |
| 1087 | TTT **A**T**T** ATT | → | TTT **-**T**-** ATT | (3) |
| 1157 | ACC A**T**T **A**TT | → | ACC A**a**T **t**TT | (3) |
| 1157 | ACC A**T**T **A**TT | → | ACC A**a**T **t**TT | (3) |
| 1173 | AA**T** T**C**A AAT | → | AA**a** T**a**A AAT | (3) |
| 1191 | GG**T** T**C**C CGT | → | GG**g** T**t**C CGT | (3) |
| 1361 | TTA A**A**T **A**TC | → | TTA AA**a**T **t**TC | (3) |

| | | | | |
|---|---|---|---|---|
| 1431 | GC**T** T**T**G AAA | → | GC**c** T**a**G AAA | (3) |
| 1483 | TTA **A**TG CCC | → | TTA **t**TG CCC | (29) |
| +1511 | GCC A**C**A TTT | → | GCC A**a**A TTT | |
| 1623 | TG**G** A**T**C TTA | → | TG**a** A**c**C TTA | (3) |
